# HAIRpred2: Human Host-Specific Prediction of Antibody-Interacting Residues Using Hybrid Physicochemical and Structural Features

**DOI:** 10.64898/2026.05.09.723672

**Authors:** Naman Kumar Mehta, Ruchir Sahni, Nishant Kumar, Gajendra P. S. Raghava

**Author notes:** Corresponding Author, Prof. Gajendra P. S. Raghava, Professor, Department of Computational Biology, Indraprastha Institute of Information Technology, Delhi, Okhla Industrial Estate, Phase III, New Delhi, India – 110020, Website: http://webs.iiitd.edu.in/raghava/. Equal contribution (Naman Kumar Mehta, Ruchir Sahni, Nishant Kumar). Mailing Address of Authors Naman Kumar Mehta (NKM), Ruchir Sahni (RS), Nishant Kumar (NK). **Author’s Biography** 1. **Naman Kumar Mehta** is currently working as Ph.D. in Computational biology from Department of Computational Biology, Indraprastha Institute of Information Technology (IIIT), New Delhi, India. 2. **Ruchir Sahni** is currently studying as an integrated BS-MS student at Indian Institute of Science Education and Research (IISER) Pune, India. He is currently working as an Intern on Project position at Department of Computational Biology, Indraprastha Institute of Information Technology (IIIT), New Delhi, India. 3. **Nishant Kumar** is currently working as Ph.D. in Computational biology from Department of Computational Biology, Indraprastha Institute of Information Technology (IIIT), New Delhi, India. 4. **Gajendra P. S. Raghava** is currently working as Professor and Head of Department of Computational Biology, Indraprastha Institute of Information Technology (IIIT), New Delhi, India.

## Abstract

1.

Prediction of conformational B-cell epitopes is critical for vaccine design, immunotherapy, and antibody engineering. To date, several host-independent computational methods have been developed for predicting antibody-interacting residues in antigen structures. However, it is well established that antigen-antibody (Ag-Ab) interactions vary depending on the host immune system indicating the importance of developing host-specific prediction models. In this study, we present, for the first time, a human host-specific method, HAIRpred2, that predicts antibody-interacting residues in an antigen from its tertiary structure. The dataset was derived from HAIRpred and comprises 277 human Ag-Ab complexes, with 221 structures used for training and 56 for independent testing. Preliminary analysis revealed that residues with a relative surface accessibility (RSA) below 0.05, corresponding to buried regions, are highly likely to be non-interacting, underscoring the importance of structural accessibility in antibody recognition. To identify the most informative features, we evaluated multiple feature representations, including RSA, large language model (LLM)-based embeddings, distance-based features, and physicochemical properties. A model trained on single-residue RSA features achieved an AUC of 0.72. Incorporating a sliding window of 15 residues to capture local structural context improved performance to an AUC of 0.75. The best performance (AUC = 0.78 on the independent test set) was achieved by integrating RSA with physicochemical descriptors. Benchmarking against existing antibody-interaction prediction methods on the same independent dataset demonstrated that HAIRpred2 outperforms current tools, further highlighting the advantage of host-specific modeling. HAIRpred2 is freely available as a web server at https://webs.iiitd.edu.in/raghava/hairpred2/.

**Highlights:**

- Development of HAIRpred2, the first human host-specific method for predicting antibody-interacting residues.
- Analysis of 277 human antigen–antibody complexes to capture host-dependent interaction patterns.
- Relative surface accessibility (RSA) identified as a key determinant, with buried residues rarely participating in interactions.
- Integration of RSA with physicochemical features achieved the best performance (AUC = 0.78) on an independent dataset.
- HAIRpred2 outperforms existing methods and is available as a web server for epitope prediction.

## 2. Introduction

The immune system is a complicated defence mechanism that works together to keep diseases from getting into the body. Antibodies are a key part of humoral immunity. They find and attach to antigens by recognising certain amino acid residues on the surface of the antigen, which are often called antibody-interacting residues. These residues together make up B-cell epitopes, which are the parts of an antigen that B-cell receptors and circulating antibodies recognise. These are important for starting an adaptive immune response. It is important to note that Ag-Ab interactions can differ among host species because of changes in immunological repertoire, germline gene utilisation, and the ways that affinity matures. Accurate identification of B-cell epitopes is crucial in vaccine production and antibody-based immunotherapy, as these areas provide the major binding sites for Ag-Ab recognition^1^.

Early computational methods for predicting B-cell epitopes mostly used propensity-based methods, like amino acid secondary structure. While these methods yielded valuable biological insights into epitope features, subsequent investigations revealed that propensity-based approaches alone do not possess adequate predictive accuracy for reliable identification^2^. This realisation led to the creation of more complex computer approaches that use machine learning algorithms and a variety of attributes derived from sequences. This restriction led to the creation of more sophisticated computational techniques that integrate machine learning algorithms and a variety of sequence-derived attributes. A lot of early predictors looked at continuous or linear B-cell epitopes. These are amino acid residues that are adjacent to each other in the antigen’s basic structure. BEPITOPE, ABCpred, and BepiPred are some of the most popular programs that leverage physicochemical features, hidden Markov models, neural networks, or propensity-based scoring algorithms to find possible epitope sites in protein sequences ^3–5^. Even though they are popular, only a tiny number of B-cell epitopes are actually continuous^6^. Furthermore, even residues that appear to form linear epitopes assume distinct three-dimensional conformations upon antibody binding, demonstrating that structural context is essential for epitope recognition beyond basic sequence alone.

Conversely, most B-cell epitopes are discontinuous, consisting of residues that are distant in primary sequence yet physically adjacent in the folded protein structure. Several structure-based predictors have been created over the years to help scientists understand how Ag-Ab interactions work. These include CEP, DiscoTope, Epitopia, PEPITO, SEPPA, Epitope3D, and SEMA, as well as other approaches shown in Table 1 ^2,7–12^. These methods incorporate structural features such as spatial proximity, solvent accessibility, and evolutionary conservation to improve conformational epitope prediction. The majority of current structure-based methods are not host-specific and are typically trained on diverse datasets that include Ag-Ab complexes from many species, despite these developments. This lack of host specificity may limit their ability to accurately identify the unique characteristics of human immune responses. This absence of host specificity may constrain their capacity to precisely delineate the distinctive characteristics of human immune responses. Structural and physicochemical examinations of antigen-antibody complexes have demonstrated that epitope regions possess unique attributes, such as relative solvent accessibility (RSA), residue depth, half-sphere exposure (HSE), electrostatic properties, hydrophobicity, isoelectric point (pI), and pKa values^8,13,14^. Recent findings indicate that residue representations derived from LLM embeddings can identify trends in the context and evolution of residues, enhancing the accuracy of predictions at the residue level.

**Table 1.**
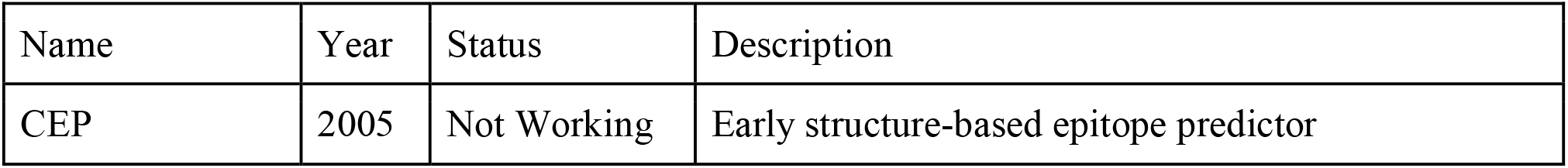
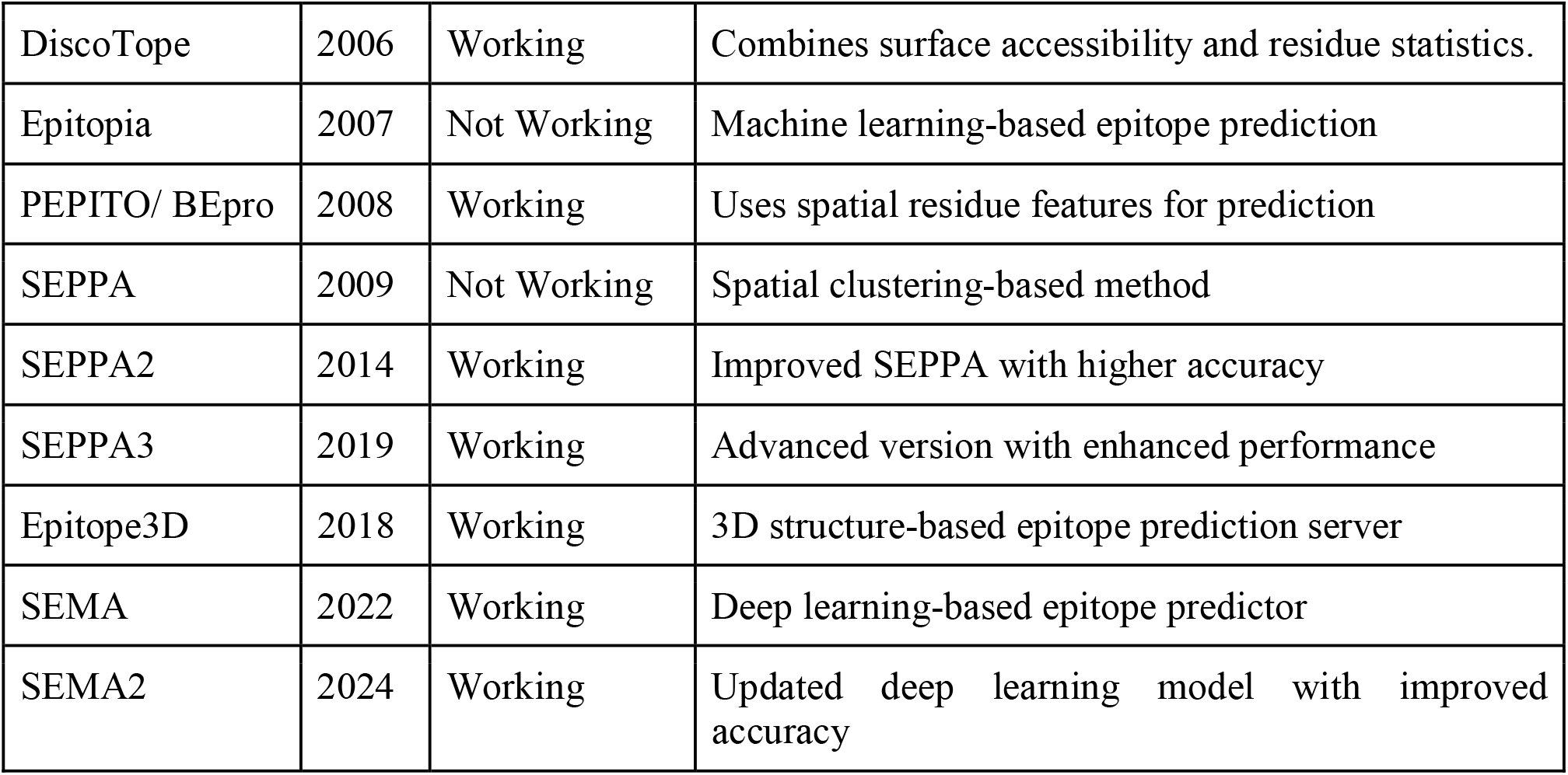
List of tools developed for the prediction of conformational or discontinuous epitopes in an antigen.

| Name | Year | Status | Description |
| --- | --- | --- | --- |
| CEP | 2005 | Not Working | Early structure-based epitope predictor |
| DiscoTope | 2006 | Working | Combines surface accessibility and residue statistics. |
| Epitopia | 2007 | Not Working | Machine learning-based epitope prediction |
| PEPITO/ BEpro | 2008 | Working | Uses spatial residue features for prediction |
| SEPPA | 2009 | Not Working | Spatial clustering-based method |
| SEPPA2 | 2014 | Working | Improved SEPPA with higher accuracy |
| SEPPA3 | 2019 | Working | Advanced version with enhanced performance |
| Epitope3D | 2018 | Working | 3D structure-based epitope prediction server |
| SEMA | 2022 | Working | Deep learning-based epitope predictor |
| SEMA2 | 2024 | Working | Updated deep learning model with improved accuracy |

In our previous research, we created HAIRpred, a computational tool that uses sequences to predict which amino acids in antigens may interact with human antibodies^15^. Although HAIRpred demonstrated useful predictive capability, its performance was inherently limited by its reliance solely on sequence-derived features. To overcome these limitations, we present HAIRpred2, an improved structure-based method for predicting human antibody-interacting residues in antigens by leveraging hybrid structural and physicochemical features derived from the antigen’s tertiary structure. The specific objectives of this study are to: (i) curate a high-quality, human host-specific Ag-Ab complex dataset; (ii) systematically evaluate and select the most informative structural and physicochemical features; (iii) develop and benchmark machine learning models for residue-level antibody-interaction prediction; and (iv) deploy HAIRpred2 as a publicly accessible web server to support the broader immunological and vaccine research community.

## 3. Results

The results section is divided into multiple groups so that we can systematically look at how alternative feature representations and learning methods help us classify Ag-Ab interacting residues. These include (i) RSA-based probability analysis to examine the relationship between solvent accessibility and residue interaction propensity, (ii) machine learning models using RSA-based pattern features, (iii) binary exposure pattern representation, (iv) distance-based RSA neighborhood features, (v) structural and physicochemical descriptors including HSE, physicochemical properties, and secondary structure features; and (vi) LLM-based sequence embeddings. Figure 1 shows the full workflow of the study, and the next subsections give further information on each analysis.

**Figure 1.**
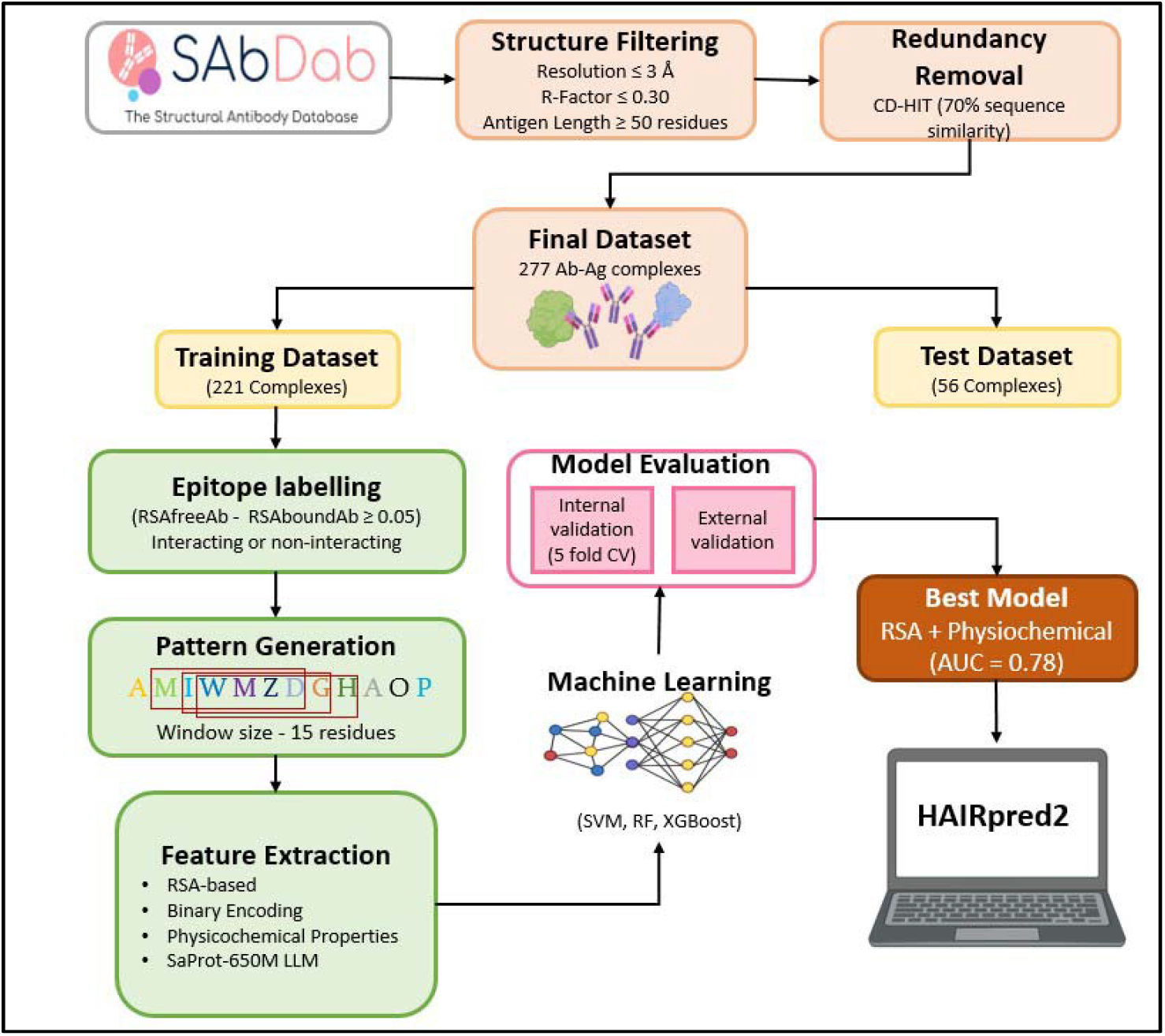
The HAIRpred2 pipeline, including all of its components.

### 3.1. Primary Analysis

The relationship between solvent accessibility and residue interaction propensity was examined using RSA-based probability analysis. Residues were categorised into predefined RSA ranges, and the probabilities of interacting and non-interacting residues were calculated for each bin to evaluate how solvent exposure influences residue behavior. The data showed a clear pattern: residues with low RSA values, which are found in buried parts of the protein structure, were very likely to not interact with other residues. On the other hand, residues with higher RSA values were more likely to be involved in interactions. As solvent accessibility increased, the probability of non-interacting residues gradually decreased while the interaction probability increased. This observation suggests that solvent-exposed residues are more accessible for intermolecular interactions, whereas buried residues primarily contribute to maintaining the internal structural stability of the protein. Supplementary Table S1 shows the precise RSA versus probability distribution of interacting and non-interacting residues, and the overall relationship across solvent accessibility ranges is illustrated in Figure 2.

**Figure 2.**
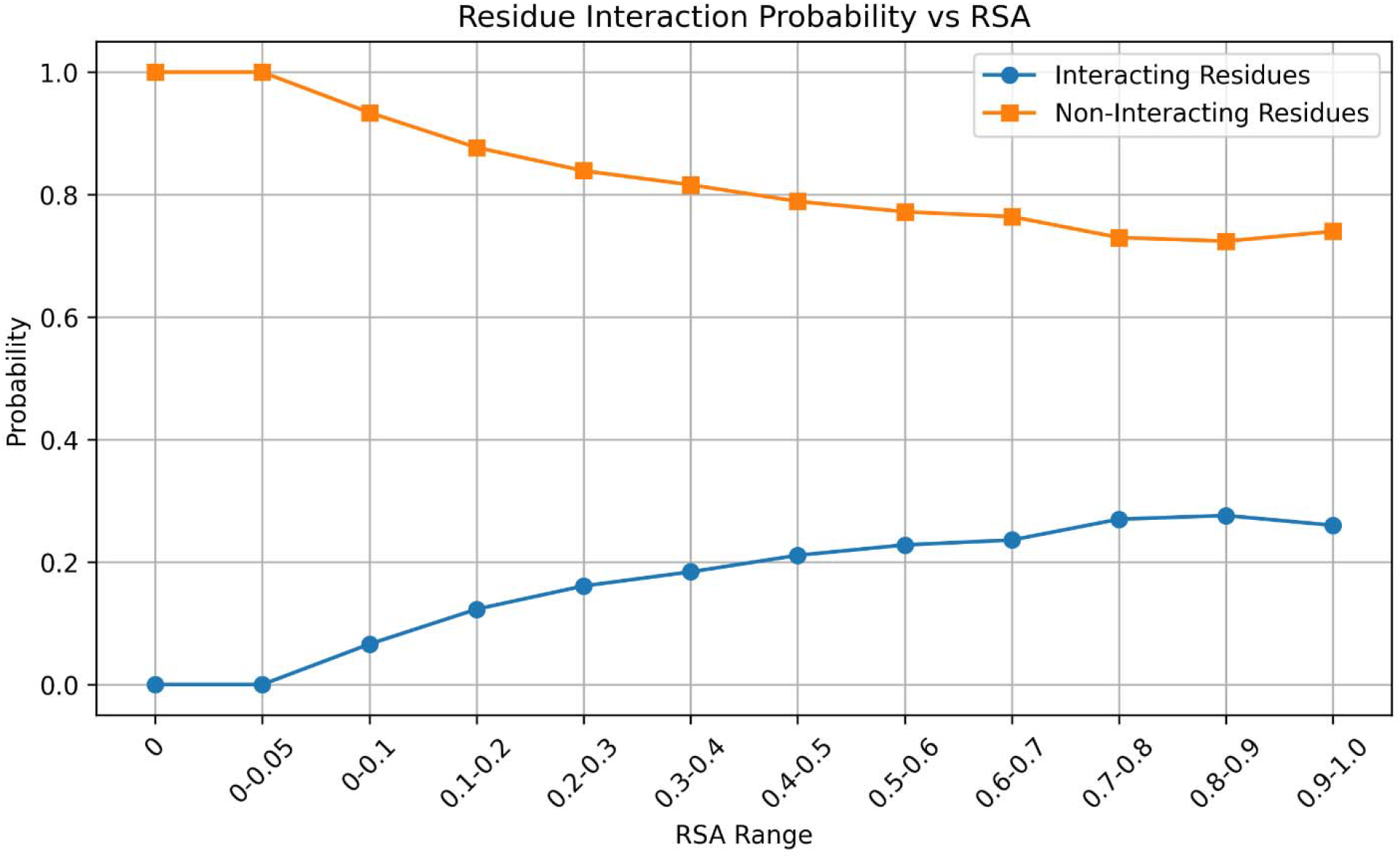
Relationship between RSA and residue interaction probability.

### 3.2. RSA single residue

The performance of AI models using single-residue RSA features was evaluated on the independent test dataset. In this representation, the RSA value of each residue was directly used as an input feature for model training. Several classifiers were trained and evaluated. The comparative performance of these models on the independent test dataset is presented in Table 2. The RSA values of individual residues are provided in Supplementary Table S2, while the complete performance metrics for both training and testing phases are available in Supplementary Table S3.

**Table 2.** Performance of machine learning models using RSA single residue.

| Model | Sensitivity | Specificity | Accuracy | MCC | AUC |
| --- | --- | --- | --- | --- | --- |
| DT | 0.6622 | 0.6248 | 0.6310 | 0.2157 | 0.7064 |
| LR | 0.6364 | 0.6477 | 0.6458 | 0.2153 | 0.7089 |
| MLP | 0.6392 | 0.6447 | 0.6438 | 0.2148 | 0.7089 |
| GNB | 0.6397 | 0.6441 | 0.6434 | 0.2148 | 0.7089 |

### 3.3. RSA Pattern Feature

The independent test dataset was used to evaluate the performance of machine learning models that use RSA-based pattern features. Specifically, a fixed sliding window of 15 residues was applied, where each window captured the solvent accessibility profile of a central residue along with its surrounding neighbors. This representation was then used to train multiple classifiers, including Extra Trees, XGBoost, Random Forest, and Logistic Regression.

Table 3 summarises the models’ comparative performance on the independent test dataset. The results show that incorporating the local solvent accessibility context through the 15-residue RSA patterns improves the ability of the models to distinguish between interacting and non-interacting residues. This highlights the importance of neighboring residue information in capturing interaction-relevant structural characteristics. Supplementary Table S4 shows the entire performance metrics for both the training and testing phases.

**Table 3.**
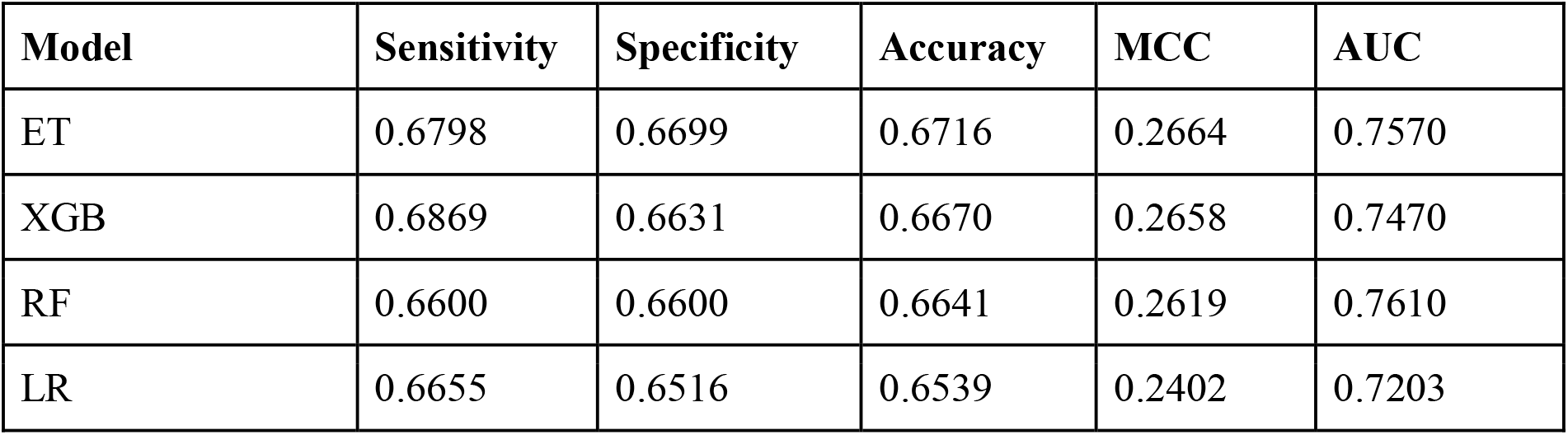
Prediction performance of machine learning models using RSA pattern representation.

### 3.4. Binary Exposure Pattern

The performance of machine learning models using the binary exposure pattern representation was evaluated on the independent test dataset. In this representation, RSA values were incorporated along with residue identity within the sliding window framework to capture the local solvent accessibility environment of antigen residues.

Several machine learning classifiers were trained using these encoded patterns to distinguish interacting and non-interacting residues. The comparative performance of the models obtained on the independent test dataset is summarized in Table 4. The results demonstrate that incorporating residue identity together with solvent accessibility information can effectively capture structural characteristics associated with residue-level interactions. Detailed performance results for both the training and testing datasets are available in Supplementary Table S5

**Table 4.** Prediction performance of machine learning models using Binary Exposure pattern representation.

| Model | Sensitivity | Specificity | Accuracy | MCC | AUC |
| --- | --- | --- | --- | --- | --- |
| ET | 0.6757 | 0.6698 | 0.6707 | 0.2621 | 0.7395 |
| XGB | 0.7029 | 0.6333 | 0.6447 | 0.2516 | 0.7295 |
| RF | 0.6914 | 0.6392 | 0.6477 | 0.2480 | 0.7300 |
| LR | 0.6557 | 0.6570 | 0.6568 | 0.2365 | 0.7200 |

### 3.5. Distance-Based RSA Feature

Distance-based RSA features were used to evaluate the predictive ability of machine learning models for residue-level Ag-Ab interactions. In this approach, spatially neighboring residues identified from the three-dimensional structure were used to capture the local structural environment of each antigen residue. We looked at two feature representations: DB-RSA, which uses the RSA values of the nearest spatial residues, and RI-RSA, which combines both residue identity and RSA information. We used an independent dataset to test models that had been trained on these attributes to tell the difference between interacting and non-interacting residues. The comparative performance of models based on DB-RSA and RI-RSA is presented in Table 5, while detailed performance metrics for both training and testing datasets are provided in Supplementary Table S6. The findings demonstrate that integrating spatial neighbourhood data with solvent accessibility offers significant structural context, hence improving the prediction of residue-level Ag-Ab interactions.

**Table 5.**
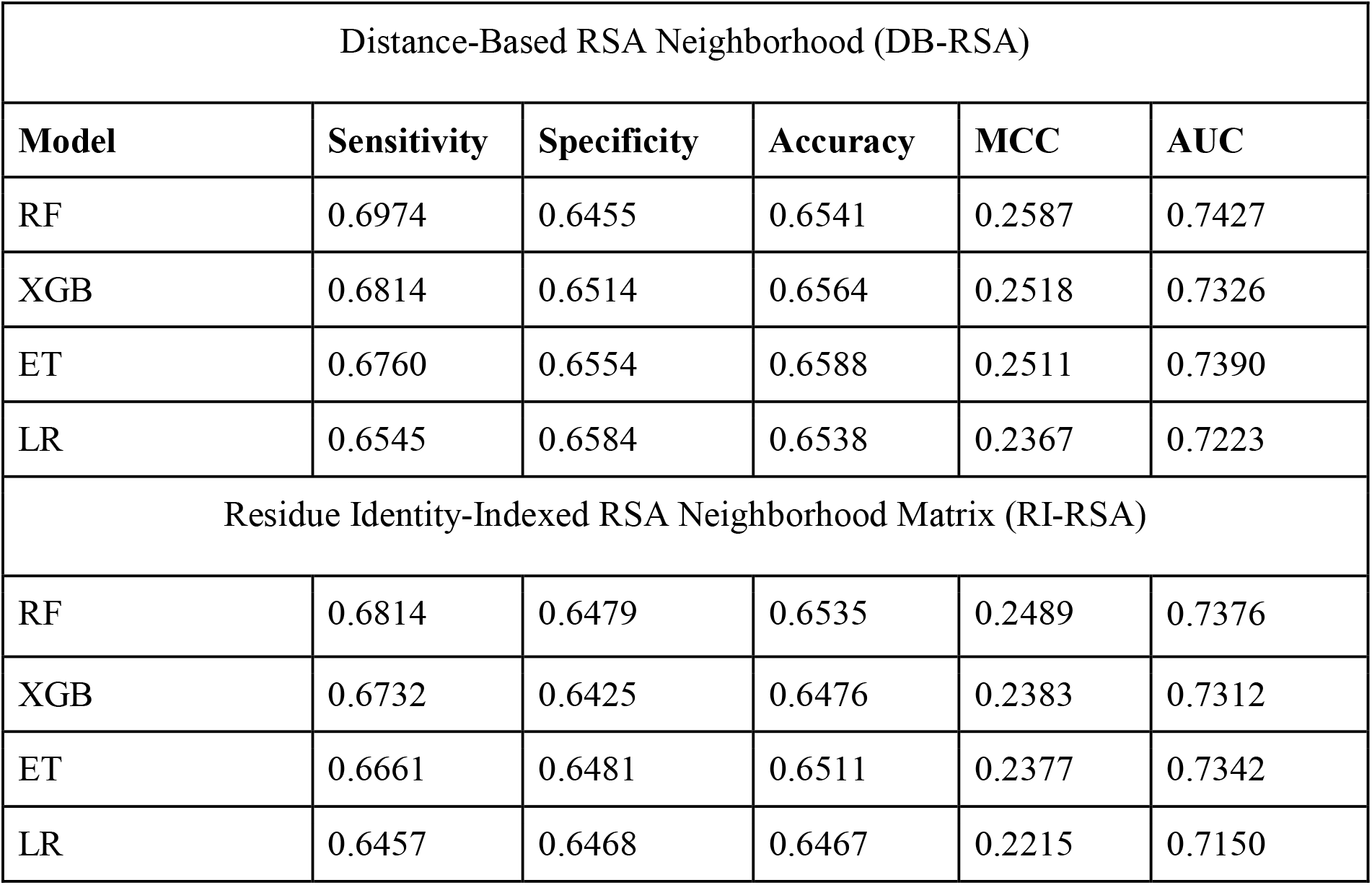
Performance of machine learning models using Distance-Based RSA Feature.

### 3.6. Half-Sphere Exposure (HSE) Features

The independent test dataset was used to assess the predictive performance of machine learning models utilising HSE-based feature representations. In this approach, HSE descriptors were used to characterise the local three-dimensional environment of each residue by quantifying the distribution of neighboring residues in the upper and lower hemispheres relative to the Cα atom. These features capture important structural aspects such as residue packing and spatial exposure, which are critical for identifying antigen–antibody interaction sites. Multiple machine learning classifiers were trained on these HSE-derived features to differentiate between interacting and non-interacting residues. The comparative results for the HSE-based models on the independent test set are presented in Table 6, while detailed performance metrics for both training and testing datasets are provided in Supplementary Table S7.

**Table 6.** Performance of machine learning models using HSE features.

| Model | Sensitivity | Specificity | Accuracy | MCC | AUC |
| --- | --- | --- | --- | --- | --- |
| ET | 0.6506 | 0.6507 | 0.6507 | 0.2284 | 0.7158 |
| RF | 0.6217 | 0.6750 | 0.6662 | 0.2276 | 0.7089 |
| XGB | 0.6228 | 0.6380 | 0.6355 | 0.1972 | 0.6891 |
| LR | 0.6334 | 0.6155 | 0.6184 | 0.1868 | 0.6703 |

### 3.7. Physicochemical Properties

To obtain both structural and biological characteristics of antigen residues, physicochemical parameters were combined with RSA-based features. For each residue, the RSA values derived from the 15-residue sliding window were combined with amino acid-specific descriptors, including isoelectric point (pI), acid dissociation constants (pKa1 and pKa2), hydrophobicity, steric parameters, and EIIP. This combined representation includes both the solvent accessibility and the inherent biological properties of the residues. This gives a full set of features for characterising epitopes.

As summarized in Table 7, interacting residues exhibited distinct physicochemical patterns compared to non-interacting ones, particularly in terms of hydrophobicity and steric properties, likely due to their preference for exposed regions on the antigen surface. Incorporating these physicochemical features alongside RSA improved model performance, increasing the AUC from 0.75 (RSA-only) to 0.78. Overall, these results indicate that combining structural exposure with biochemical attributes strengthens the model’s ability to distinguish interacting from non-interacting residues. Additional details are available in Supplementary Table S8.

**Table 7.** Performance of machine learning models using RSA with Physicochemical Properties.

| Model | Sensitivity | Specificity | Accuracy | MCC | AUC |
| --- | --- | --- | --- | --- | --- |
| RF | 0.7193 | 0.6544 | 0.6651 | 0.2823 | 0.7800 |
| ET | 0.7270 | 0.6548 | 0.6667 | 0.2883 | 0.7705 |
| XGB | 0.7034 | 0.6573 | 0.6649 | 0.2730 | 0.7584 |
| LR | 0.6710 | 0.6532 | 0.6562 | 0.2456 | 0.7238 |

### 3.8. Secondary Structure Features

We used DSSP to get secondary structure features and then coupled them with the RSA-based sliding window to get the local structural context of each residue. Table 8 shows that residues that interact with each other were more likely to be discovered in coil and loop areas, while residues that don’t interact with each other were more likely to be found in helices or β-sheets. Adding secondary structure information alongside RSA improved the model’s ability to discriminate interacting from non-interacting residues, highlighting the importance of local backbone conformation in Ag-Ab interactions. Further detailed performance analyses are presented in Supplementary Table S9.

**Table 8.**
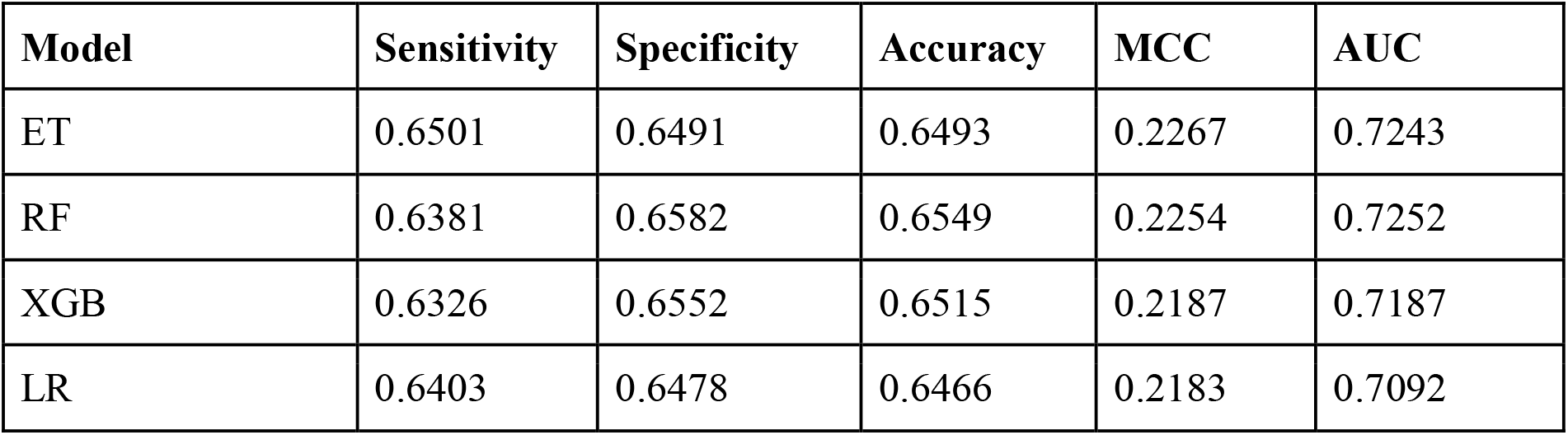
ML results for secondary structure-based residue classification.

### 3.9. Large Language Model (LLM)-Based Features

The SaProt-650M model was used in an end-to-end prediction to see how well LLM-derived sequence representations worked. We evaluated both pretrained and fine-tuned versions of the model to see if adapting it to a specific job enhances its ability to make predictions.

The pretrained model demonstrated its intrinsic capacity to extract substantial biochemical and structural information from protein sequences by establishing a robust baseline. However, the model’s consistent improvements in all of the assessment metrics were observed when the model was fine-tuned on the Ag-Ab interaction dataset. This indicates that the model acquired a greater understanding of the characteristics that are critical for interactions. The fine-tuned model was more effective in distinguishing between interacting and non-interacting residues, demonstrating the effectiveness of task-specific training. The comparison between the pretrained and fine-tuned SaProt-650M models is illustrated in Table 9.

**Table 9.** Performance comparison of pretrained and fine-tuned SaProt-650M models.

| Model Variant | Sensitivity | Specificity | Accuracy | MCC | AUC |
| --- | --- | --- | --- | --- | --- |
| Pretrained SaProt-650M | 0.511 | 0.510 | 0.510 | 0.016 | 0.514 |
| Fine-tuned SaProt-650M | 0.608 | 0.610 | 0.610 | 0.164 | 0.654 |

After testing the end-to-end predictive performance, residue-level embeddings were taken from both the pretrained and fine-tuned versions of the SaProt model to see how well they worked as input features for downstream ML-based interaction prediction. The performance of machine learning models trained using fine-tuned SaProt embeddings as input features is summarized in Table 10. The results shown in the table illustrate the comparative effectiveness of embedding-based feature representations for downstream classification of antibody-interacting residues.

**Table 10.** Performance of ML models on SaProt-650M fine-tuned models embeddings.

| Model | Sensitivity | Specificity | Accuracy | MCC | AUC |
| --- | --- | --- | --- | --- | --- |
| GNB | 0.600 | 0.517 | 0.531 | 0.087 | 0.610 |
| XGB | 0.656 | 0.477 | 0.506 | 0.099 | 0.594 |
| MLP | 0.670 | 0.443 | 0.480 | 0.085 | 0.591 |

### 3.10. Benchmarking

To achieve an objective assessment, HAIRpred2 was benchmarked against well-known structure-based epitope prediction tools, such as Sema, SEPPA 3.0, ElliPro, epitope3D, and DiscoTope-2.0, using our test dataset ^11,12,16–18^. Predictions from all methodologies were validated as labels and assessed using sensitivity, specificity, accuracy, MCC and AUC. The benchmarking results indicate notable differences in prediction behavior among the compared tools. For instance, Sema demonstrated very high specificity but extremely low sensitivity, suggesting a strong bias toward predicting non-interacting residues. Likewise, SEPPA 3.0 exhibited considerable sensitivity, although it demonstrated relatively low specificity and MCC, indicating restricted balanced differentiation between interacting and non-interacting residues.

In contrast, HAIRpred2 achieved a more balanced performance across sensitivity and specificity, resulting in improved MCC and AUC values. This balanced discrimination indicates that HAIRpred2 is better able to identify true interacting residues without excessively increasing false positives. The findings in Table 11 indicate that the structure-integrated hybrid framework of HAIRpred2 exhibits superior performance compared to current structure-based epitope prediction tools on the independent validation dataset.

**Table 11.** Benchmarking of different models on our validation data.

| Method | Sensitivity | Specificity | Accuracy | MCC | AUC |
| --- | --- | --- | --- | --- | --- |
| Sema2 | 0.029 | 0.978 | 0.831 | 0.017 | 0.503 |
| SEPPA 3.0 | 0.667 | 0.390 | 0.436 | 0.044 | 0.528 |
| ElliPro | 0.347 | 0.699 | 0.641 | 0.038 | 0.523 |
| Epitope3D | 0.560 | 0.521 | 0.540 | 0.071 | 0.531 |

## 4. Discussion and Conclusion

Antibodies are important parts of the adaptive immune system that can recognise foreign antigens by looking for certain surface-exposed areas called epitopes. Precise identification of antibody-interacting residues is essential for comprehending antigen-antibody recognition and for enhancing vaccine research, therapeutic antibody creation, and diagnostic applications. In our previous work, HAIRpred, we tackled many shortcomings of generalised epitope prediction methods by concentrating solely on empirically confirmed human antigen-antibody complexes. By combining sequence-derived, evolutionary, and structural features, most notably RSA and position-specific scoring matrix (PSSM) profiles, HAIRpred achieved a strong performance of AUC of 0.72 on independent datasets and demonstrated the importance of host specificity in improving predictive accuracy. However, HAIRpred primarily relied on sequence-driven representations and limited structural descriptors, which constrained its ability to fully capture the 3D context and spatial proximity that govern antibody binding.

To address these limitations, we created HAIRpred2, an enhanced structure-based framework utilising the same curated dataset of 277 high-quality human antigen-antibody complexes obtained from experimentally resolved structures. Unlike its predecessor, HAIRpred2 explicitly incorporates detailed structural information from antigen 3D conformations. Residue-level labels were defined based on changes in RSA between free and antibody-bound states, enabling precise identification of direct interface residues. We comprehensively assessed various feature categories, encompassing RSA-based representations, half-sphere exposure, secondary structure, and essential physicochemical features such as pI, pKa1, pKa2, hydrophobicity, steric parameters, and EIIP. In addition, we explored embeddings derived from a large language model to assess whether contextual sequence representations further enhance prediction.

Our findings indicate that structural descriptors substantially outperform sequence-only representations for residue-level interaction prediction. Among all tested configurations, a hybrid feature set combining RSA with selected physicochemical properties achieved the best performance, reaching an AUC of 0.78 on the independent test dataset, an appreciable improvement over HAIRpred. While fine-tuned language model embeddings improved over pretrained variants, they did not surpass the predictive power of structure-informed hybrid features, underscoring the critical importance of spatial exposure and biochemical complementarity in Ag-Ab recognition. Comparative benchmarking against established structure-based epitope prediction tools further confirmed the robustness and balanced performance of HAIRpred2.

In summary, HAIRpred2 represents a clear progression from the original HAIRpred framework, shifting from a largely sequence-focused, host-specific approach to a comprehensive structure-based prediction model. By integrating detailed structural information with complementary physicochemical features, HAIRpred2 achieves improved predictive accuracy and finer residue-level resolution. This work reinforces the critical role of structural context in Ag-Ab interactions and provides a reliable and accessible tool for immunological studies, rational vaccine design, and therapeutic antibody development.

## 5. Methods

### 5.1. Dataset collection

In the current study, we have extracted the Ag-Ab structural complexes from the SAdDab database^19^ for the human host only. The dataset used in this study was adopted from HAIRpred, where Ag-Ab complexes were curated using stringent quality criteria. Only structures with a resolution less than 3 Å and an R-factor of 0.30 have been selected for our study. The antigens in a complex that have a length of less than 50 amino acids were removed from the dataset. We have used CD-HIT software with the standards of 70% sequence identity to take care of the redundancy ^20^. Eventually, we were left with 277 Ag-Ab complexes that fulfill our dataset selection criteria. Our final dataset has 221 training and 56 testing complexes, which is available at https://webs.iiitd.edu.in/raghava/hairpred2/download.

### 5.2. Epitope labeling

Residue-level epitope annotations were generated based on the change in RSA between the free antigen structure and the antigen structure bound to the antibody. RSA values were computed using DSSP. A residue was considered an interacting epitope residue if it satisfied the following condition:

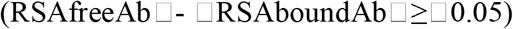

This criterion reflects the reduction in solvent accessibility when the antigen interacts with the antibody. Residues showing a significant decrease in RSA upon binding were labeled as interacting residues, whereas the remaining residues were labeled as non-interacting residues.

### 5.3. Primary Analysis

To examine the relationship between solvent accessibility and residue interaction propensity, a residue-level analysis based on RSA was performed. Residues labeled as non-interacting were grouped according to their RSA values, and the RSA values were divided into several bins representing different levels of solvent exposure. The probability of non-interacting residues was determined for each RSA group by calculating the fraction of non-interacting residues within each group, as well as for the combined residue set, for all twenty amino acid types. The calculated probabilities were subsequently used to generate RSA-based probability diagrams to demonstrate the correlation between the probability of residues not interacting and solvent accessibility, as well as the inverse. This analysis was conducted to ascertain the behaviour of residues at varying levels of exposure and to aid in the development of structural elements that will be used in subsequent modelling processes.

### 5.4. Epitope Pattern Generation

A sliding window-based pattern generating approach was used to show how the interaction behaviour of each antigen residue is affected by nearby residues. A fixed-length window of 15 residues was established for each residue in the antigen sequence. This window included the core residue and its neighbouring residues on both sides. If the window extended beyond the N or C-terminal ends of the sequence, the missing positions were filled using padding values to maintain a consistent pattern length. This method made it possible to show the local sequence and structural context around each residue in the same way. The patterns that were made with the centred residue were then employed for feature encoding and model construction.

### 5.5. RSA single residue

RSA was used as a primary structural feature to represent the solvent exposure of antigen residues. The RSA for each residue was computed using DSSP from the 3D structures of Ag-Ab complexes. The RSA value of each residue was explicitly utilised as an input feature for the machine learning model in the single-residue representation. The RSA values range from 0 to 1, with lower values signifying buried residues and higher values denoting solvent-exposed residues. RSA is a critical discriminatory feature for the identification of potential epitope residues, as antibody binding primarily occurs on exposed regions of the antigen. Additional feature representations were constructed using these values to capture the local solvent accessibility environment and neighbouring residue context for machine learning-based classification.

### 5.6. RSA Pattern Feature

Using the already established sliding window method, we created RSA-based patterns to get the solvent accessibility profile of residues and their surrounding environment. A 15-residue window was created for each antigen residue, and the RSA values of all the residues in the window were used as feature inputs. This picture keeps the background of the core residue’s local solvent accessibility. The interaction labels for the core residues (either interacting or not) were already set during the epitope labelling phase. Machine learning models used the RSA-based patterns and their labels as input features. The models were trained to predict if the centre residue in each pattern is an interacting or non-interacting residue.

### 5.7. Binary Exposure Pattern

Binary exposure patterns were created to represent residue-specific solvent accessibility within the sliding window framework. For each antigen residue, a window of 15 neighboring residues was considered, and features were organised according to the 20 standard amino acid types. In this representation, the RSA value of each residue was assigned to the position corresponding to its amino acid type, while all other positions were filled with zero. This encoding preserves both residue identity and solvent accessibility information within the local neighborhood of the central residue. The interaction label assigned during the epitope labeling step was retained for the central residue, and the resulting encoded patterns were used as input features for machine learning models to classify residues as interacting or non-interacting.

### 5.8. Distance-Based RSA Feature

We created Distance-Based RSA Neighbourhood (DB-RSA) characteristics to use the three-dimensional coordinates of antigen residues to add information about structural proximity. Coordinates of the Cα atom were taken from antigen structures, and Euclidean distances were used to find spatial neighbours. For each antigen residue, the 15 nearest residues in three-dimensional space were chosen to make a structural neighbourhood, no matter where they were in the sequence. The RSA values of these neighbouring residues were utilised to make a feature vector that shows how accessible the solvent is surrounding the core residue. The interaction label given during the epitope labelling step was kept for the central residue.

We also constructed a Residue Identity-Indexed RSA Neighbourhood Matrix (RI-RSA) using the same spatial neighbourhood. In this encoding, the RSA value of each neighbouring residue was given based on its amino acid type, while the other locations were filled with zeros. This kept both the identity of the residue and the information about how accessible it is to solvent. These encoded patterns were used as input features for machine learning models to classify residues as interacting or non-interacting.

### 5.9. Half-Sphere Exposure Features

We used the three-dimensional structure of the antigen to figure out HSE characteristics, which show the local structural environment around antigen residues. HSE counts how many residues are next to a central residue in two hemispheres that are determined in relation to the Cα-Cβ vector. These hemispheres reflect the upper and lower spatial areas around the residue. These features tell us about the packing density in the area and how the neighbouring residues are spread out in space. This helps us tell the difference between areas that are exposed to the surface and areas that are more tightly packed with structure. We used the computed HSE values as structural characteristics and fed them into machine learning algorithms to classify residues as interacting or non-interacting.

### 5.10. Physicochemical Properties

Physicochemical properties were combined with RSA-based features to integrate biochemical characteristics of residues along with structural exposure information. For each antigen residue, RSA values from the 15-residue sliding window were combined with a number of amino acid descriptors, including isoelectric point (pI), acid dissociation constants (pKa1 and pKa2), hydrophobicity, steric parameters, and electron-ion interaction potential (EIIP). The amino acid type in the pattern determined these attributes, which let the feature representation show both the biological and solvent accessibility of residues. We used the combined data as input for machine learning models that could classify the difference between residues that were interacting or non-interacting.

### 5.11. Secondary Structure

We used DSSP to get secondary structure features from antigen structures to capture the local conformation of antigen residues. Each residue was assigned a secondary structure state based on its backbone hydrogen bonding pattern and geometric configuration. These states were encoded and combined with the RSA-based sliding window features to represent the structural context of residues and their neighbouring positions, indicating whether residues are located in helix, sheet, or coil regions. The resulting combined features were used as input for machine learning models to classify residues as interacting or non-interacting.

### 5.12. Large Language Model (LLM)-Based Features

Protein language model-based embeddings were used to capture contextual information from antigen sequences. In this study, residue-level embeddings were generated using the SaProt-650M protein language model, which integrates both sequence and structural information^21^. Structure-informed antigen sequences were provided as input to the pretrained SaProt model to obtain contextual embeddings for each residue. To better adapt the model to the epitope prediction task, the SaProt-650M model was further fine-tuned using the training dataset of antigen residues and their interaction labels. The resulting residue-level embeddings were then extracted and used as feature representations. These embeddings were further used as input features for machine learning models to predict residues as interacting or non-interacting.

### 5.13. Machine learning models

We explored several supervised machine learning approaches to evaluate the effectiveness of the proposed residue-level features, including support vector machines, random forests, and gradient boosting-based classifiers. To ensure the models were reliable and could generalise well to unseen data, we implemented a five-fold cross-validation scheme on the training dataset.

Five-fold cross-validation was used on the training dataset to make sure that the model construction was accurate. We divided the data into five groups. We used one group for validation and the other four groups for training. GridSearchCV was used to systematically find the optimum parameter settings for each model and feature combination by optimising hyperparameters. The refined models were then retrained on the entire training dataset and assessed on a separate test set to evaluate their generalisation capability. The evaluation of model performance was conducted by sensitivity, specificity, accuracy, and Matthews correlation coefficient (MCC), as detailed below.

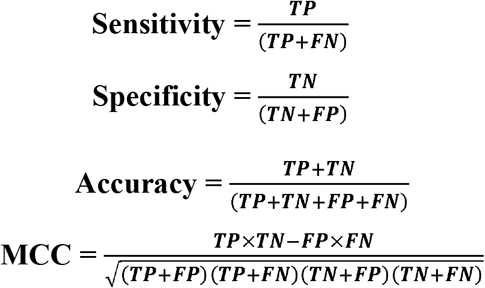

### 5.14. Web server

We created an upgraded web server, HAIRpred2, to facilitate researchers in predicting antibody-interacting residues in antigens. The server offers an intuitive interface constructed using HTML, JavaScript, and PHP, facilitating the rapid submission and analysis of antigen structures. Users may upload protein structures in conventional formats, and the service provides residue-level predictions along with downloadable results. Besides the web-based implementation, HAIRpred2 is offered as a standalone program and as a Python module that can be installed using pip.

## Abbreviations

Ag-Ab: Antigen-Antibody
AUC: Area Under the Curve
DB-RSA: Distance-Based Relative Solvent Accessibility
DT: Decision Tree
EIIP: Electron-Ion Interaction Potential
ET: Extra Trees (Extremely Randomized Trees)
GNB: Gaussian Naive Bayes
HSE: Half-Sphere Exposure
LLM: Large Language Model
LR: Logistic Regression
MCC: Matthews Correlation Coefficient
ML: Machine Learning
MLP: Multilayer Perceptron
pI: Isoelectric Point
pKa: Acid Dissociation Constant
PSSM: Position-Specific Scoring Matrix
RF: Random Forest
RI-RSA: Residue Identity-Indexed-Relative Solvent Accessibility
RSA: Relative Solvent Accessibility
XGB: Extreme Gradient Boosting

## Author’s Contribution

**Naman Kumar Mehta (NKM):** Methodology, Validation, Formal analysis, Software and Coding, Webserver implementation, writing original draft, Reviewing and editing the original draft. **Ruchir Sahni (RS):** Collected and Proceed the datasets, Conceptualization, Methodology, Formal analysis, Write Code, Data Curation, Reviewing and editing the original draft. **Nishant Kumar (NK):** Conceptualization, Methodology, Formal analysis, Data Curation, Reviewing and editing the original draft. **Gajendra Pal Singh Raghava (GPSR):** Supervision, Conceptualization, project administration, Reviewing and editing the original draft, Investigation, Validation.

## Data Availability statement

The dataset generated for this study can be accessed on the ‘hairpred2’ web server at http://github.com/raghavagps/hairpred2. The source code of this study is publicly available on Github and can be found http://github.com/raghavagps/hairpred2.

## Acknowledgement

The authors express their gratitude to the University Grants Commission (UGC) for the generous fellowship and financial support, they also Thank the Department of Computational Biology, IIITD, New Delhi, for the excellent infrastructure and facilities. The authors would like to acknowledge the Department of Biotechnology (DBT) for the infrastructure grant awarded to the Institute. Furthermore, they would like to acknowledge Biorender.com for the creation of the figures utilized in this work.

